# Barcoding the Past: A Cost-Effective ONT-Based Workflow for COI Recovery from Century-Old Spider Museum Specimens

**DOI:** 10.64898/2026.11.20.762721

**Authors:** Łukasz Trębicki, Marko Prous, Monika Trębicka, Mirosława Dabert, Marko Mutanen

## Abstract

Natural history collections contain vast amounts of undocumented biodiversity, yet degraded DNA has long limited their integration into modern barcode reference systems. Here, we present a cost-effective Oxford Nanopore Technologies (ONT)-based workflow for recovering COI barcodes from historical spider museum specimens, using the taxonomically challenging jumping spider genus *Cytaea* as a model system. We developed a mini-barcoding approach based on six short overlapping COI fragments (∼130 bp), amplified in two multiplex PCR reactions and sequenced on ONT Flongle flow cells. The workflow was applied to 288 museum specimens ranging from 13 to 141 years old, preserved primarily in 75% ethanol under standard museum conditions. COI data were recovered from 284 of 288 specimens (98.6%). Reconstructed sequences ranged from 115 to 663 bp, and 260 of the 284 sequences (91.5%) contained at least 300 bp of resolved sequence. Amplification success differed strongly among primer pairs (31.6–98.3%), highlighting the importance of marker optimization for degraded DNA. Although older specimens generally yielded shorter sequences, informative barcode data were recovered even from century-old material. The combined ONT and Sanger dataset substantially expanded the documented barcode diversity of *Cytaea*, revealing 54 BINs, including 53 newly added to BOLD. Because fragmented mini-barcodes complicate standard clustering approaches, we placed short sequences onto a reference backbone constructed from full-length COI barcodes. This enabled their assignment to major lineages, although placement confidence varied among specimens. Our results demonstrate that historical museum specimens can be efficiently transformed into usable barcode resources using a simple, scalable, and affordable ONT-based workflow. This approach provides a practical bridge between legacy museum collections and modern DNA-based taxonomy, enabling degraded historical material to be incorporated into global biodiversity reference systems.

## Introduction

Biodiversity is declining at an unprecedented rate, with species disappearing faster than they are formally described (Cardinale *et al*., 2012; Cowie, Bouchet and Fontaine, 2022). As a result, documenting global biodiversity has become an urgent priority, particularly in the context of the ongoing taxonomic impediment (Mittelbach *et al*., 2007; Mora *et al*., 2011). Despite over 250 years of Linnaean taxonomy (Linnaeus, 1758; Manktelow, 2010) and more than one million described species (Mora *et al*., 2011), the vast majority of biodiversity remains undocumented (Mittelbach *et al*., 2007). This gap is especially pronounced in tropical regions, which harbour the highest levels of diversity yet remain understudied and highly threatened (Raven *et al*., 2020).

The situation is further exacerbated by a shortage of taxonomic expertise, limited funding, and declining recognition of the discipline (Engel *et al*., 2021). Traditional morphology-based taxonomy remains fundamental; however, it is increasingly constrained by time-intensive workflows, limited access to specimens, and the presence of morphologically cryptic species (Bickford *et al*., 2007; Pfenninger and Schwenk, 2007; Fišer, Robinson and Malard, 2018; Britz, Hundsdörfer and Fritz, 2020; Li and Wiens, 2023; Braby, Hsu and Lamas, 2024). To address these limitations, molecular approaches have become a key element of modern taxonomy.

Among them, DNA barcoding, based on a ∼650 bp fragment of the mitochondrial cytochrome c oxidase subunit I (COI) gene (Hebert *et al*., 2003), is widely adopted for animal identification. It enables rapid species identification, including immature specimens, and large-scale biodiversity assessments, supported by reference systems such as BINs and MOTUs (Hebert and Gregory, 2005; Ratnasingham and Hebert, 2013; Hebert *et al*., 2016; Hobern, 2021; Ratnasingham *et al*., 2024; NCBI, 2025). Although COI barcoding has proven highly effective in a wide range of studies, its application to degraded DNA remains challenging, limiting its use for historical specimens. This represents a major limitation, as natural history collections house vast numbers of specimens, including type material and samples from regions that are no longer accessible or habitats that have been lost or transformed (Suarez and Tsutsui, 2004; Letsch *et al*., 2025; Kapli *et al*., 2026). Advances in museomics, which applies modern molecular techniques to historical specimens, have opened new opportunities to recover genetic information from degraded DNA preserved in these collections (Raxworthy and Smith, 2021; Levesque-Beaudin *et al*., 2023; Kapun *et al*., 2025). However, DNA extracted from such material is typically highly fragmented, present in low concentrations, and difficult to amplify and sequence using standard protocols (Miller *et al*., 2013; Staats *et al*., 2013; Derkarabetian, Benavides and Giribet, 2019; Gauthier *et al*., 2020; Raxworthy and Smith, 2021; Ferrari *et al*., 2023; Todisco *et al*., 2024). Fragmentation of historical DNA strongly constrains the recovery of long amplicons, making short targeted regions more suitable for sequencing of older specimens (Ferrari *et al*., 2023). Mini-barcoding approaches, targeting short COI fragments (∼100–300 bp), provide a practical alternative for degraded DNA and have been successfully implemented using high-throughput platforms (Hajibabaei *et al*., 2006; Prosser *et al*., 2016). However, these approaches are often costly and require specialised infrastructure. Recent advances in third-generation sequencing, particularly Oxford Nanopore Technologies (ONT), offer a more accessible and scalable solution. Portable nanopore sequencers such as MinION, which can also accommodate lower-capacity Flongle flow cells via an adapter, enable cost-effective sequencing of large numbers of specimens and facilitate the discovery of previously undescribed biodiversity, including so-called “dark taxa” (Page, 2016; Srivathsan *et al*., 2019, 2024; Hartop *et al*., 2022, 2024; Hebert *et al*., 2025; Meier *et al*., 2025; Meier, Lawniczak and Srivathsan, 2025). Although COI mini-barcodes can be useful for identifying specimens with degraded DNA, they may provide lower species-level resolution than full-length barcodes (Hajibabaei *et al*., 2006; Meusnier *et al*., 2008). To address this issue, phylogenetic placement methods such as EPA-ng allow short sequences to be probabilistically assigned to branches of a reference tree constructed from full-length barcodes (Barbera *et al*., 2019). This approach enables the integration of heterogeneous datasets and provides a robust framework for interpreting short-read data in a phylogenetic context (Berger, Krompass and Stamatakis, 2011; Czech, Barbera and Stamatakis, 2019). These methodological advances are particularly valuable for taxonomic studies of poorly known and highly diverse groups, where species identification and delimitation are often complicated by limited morphological documentation and the scarcity of fresh material. In such cases, historical museum collections provide an essential source of specimens that can be linked to newly generated molecular data, helping to bridge gaps between traditional taxonomy and DNA-based species identification.

These approaches are particularly relevant to taxonomically challenging groups such as the jumping spider genus *Cytaea* Keyserling, 1882, distributed across the Oriental and Australian regions (Richardson *et al*., 2006; Żabka, Patoleta and Trębicki, 2019). The genus currently includes 44 nominal species (World Spider Catalog, 2026), many of which are poorly documented, lacking adequate illustrations or descriptions of both sexes. Morphological diversity within the genus raises questions about its monophyly, and species delimitation is further complicated by subtle morphological variation. Recent revisionary work based on museum material has led to re-examination of type specimens (Patoleta and Trębicki, 2015; Trębicki, Patoleta and Gardzińska, 2016), taxonomic reassignments, and the description of new taxa (Trębicki, Patoleta and Marusik, 2025), including the redescription of the type species *C. alburna* Keyserling, 1882 supported by COI data (Trębicki *et al*., 2021). Despite these advances, the diversity of *Cytaea* remains substantially underestimated. Examination of more than 1,300 museum specimens revealed a high number of undescribed taxa, while the limited availability of fresh material restricts the use of standard molecular approaches. This highlights the need for methods capable of recovering genetic data from historical specimens and integrating them into a phylogenetic framework. Museum genomics requires methodological frameworks specifically adapted to degraded DNA rather than direct transfer of protocols developed for fresh material (Ferrari *et al*., 2023). In this study, we develop a cost-effective workflow for COI mini-barcoding of museum specimens using Oxford Nanopore Technologies. By amplifying six short, overlapping COI fragments (∼130 bp) in a multiplex PCR approach and sequencing them on the ONT Flongle platform, we recover DNA barcodes from specimens up to 141 years old. We further integrate these data using phylogenetic placement methods, enabling the incorporation of short-read sequences into a reference backbone. This approach expands geographic and taxonomic sampling and provides a scalable framework for accelerating biodiversity research using museum collections.

## Material and Methods

### Specimen sampling and metadata

This study is based on specimens deposited in several natural history museum collections. The material originated from various biodiversity surveys conducted across Australasia. Most specimens were preserved in 75% EtOH, with a small number (n = 5) from the Museum and Art Gallery of the Northern Territory and the Western Australian Museum preserved in 96% EtOH. All specimens were stored at room temperature prior to analysis. The specimens were sourced from the following institutions: Australian Museum, Sydney (AMS); American Museum of Natural History, New York (AMNH); National Museum of Natural History (Naturalis), Leiden, The Netherlands (NMNH); Queensland Museum, Brisbane (QMB); Upper Silesian Museum in Bytom (USMB); Museum and Art Gallery of the Northern Territory (MAGNT); Zoological Museum, University of Turku (ZMUT); Western Australian Museum, Perth (WAMP); Museum and Institute of Zoology, Polish Academy of Sciences, Warsaw (MIZ PAS); Bernice Pauahi Bishop Museum, Honolulu (BPBM); Lee Kong Chian Natural History Museum, National University of Singapore (LKCNHM); and University of Łódź, Department of Invertebrate Zoology and Hydrobiology (UNILODZ). Specimen numbers and sequencing approaches are summarised by collection in Table 1.

**Table 1.**
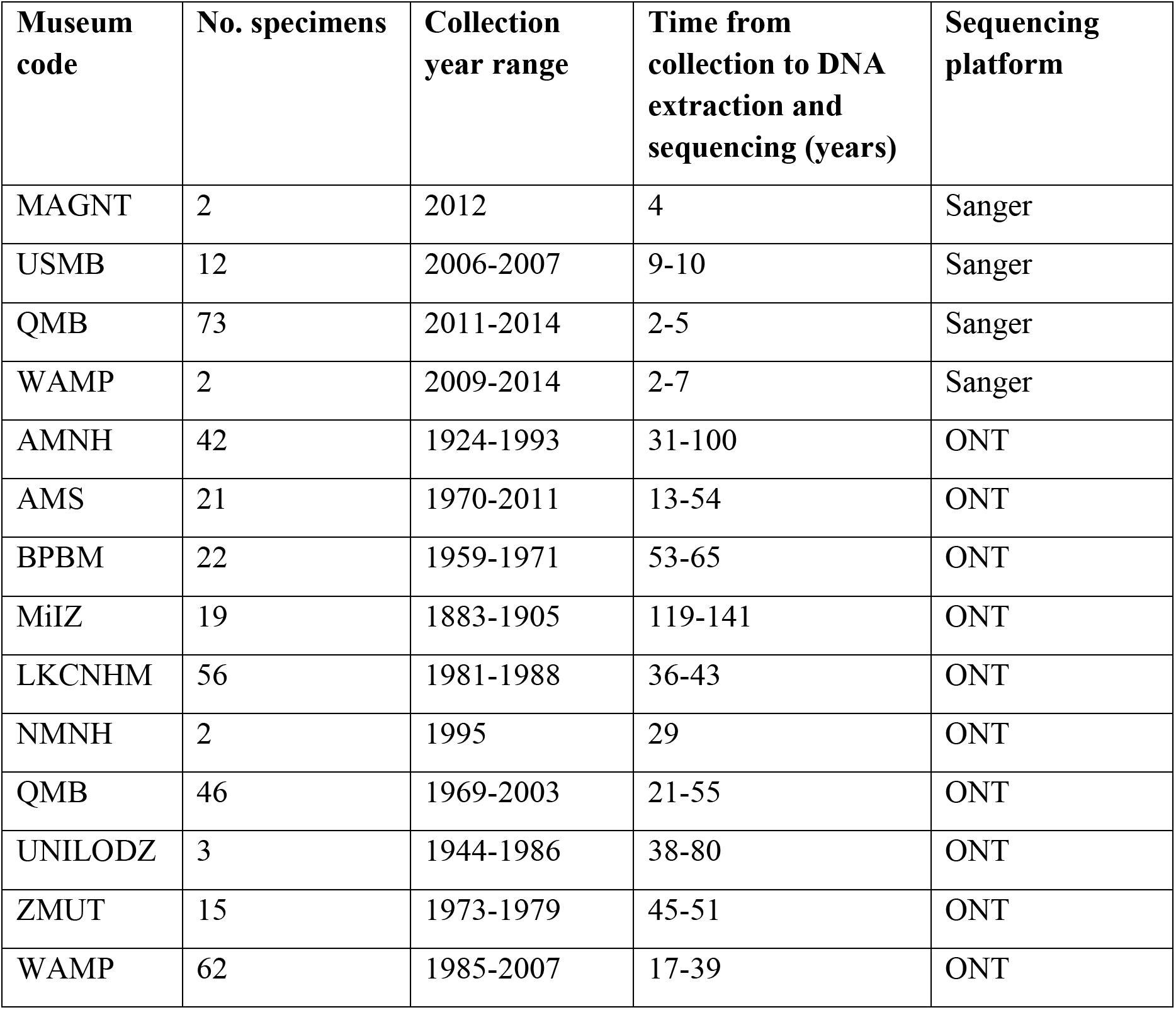
Summary of sampled museum collections, including specimen numbers, collection year ranges, time from collection to sequencing, and sequencing platform.

### Morphological examination

Morphological examinations were conducted using Nikon SMZ800, SMZ1000, and Nikon Ci compound microscopes. The analysed material spans a broad spectrum of diversity within the genus *Cytaea* and consists predominantly of museum specimens, a substantial proportion of which likely represent undescribed taxa. Specimens were partitioned into provisional morphologically delimited units (hereafter morphospecies) based on comparative morphological examination, including reference to type material of described species of *Cytaea*. These units were not intended as formal taxonomic hypotheses but were used as operational entities for downstream analyses. The morphospecies were subsequently aggregated into informal species groups according to their overall morphological affinity to described taxa, providing a higher-level framework of phenotypic similarity. This framework was used as an external reference in maximum likelihood (ML) phylogenetic analyses and phylogenetic placement procedures, allowing an independent assessment of topological congruence and the reliability of sequence assignments obtained with the tested protocol. The provisional species groups were defined as follows: the alburna group, including *C. alburna* Keyserling, 1882 (type species of the genus) and *C. aspera* (Karsch, 1878); the frontaligera group, comprising *C. frontaligera* (Thorell, 1881); the mitellata group, including *C. mitellata* (Thorell, 1881) and *C. plumbeiventris* (Keyserling, 1881); the sinuata group, including *C. sinuata* (Doleschall, 1859) and *C. nimbata* (Thorell, 1881); and a set of ungrouped taxa treated as varia. Specimens included in the phylogenetic analysis are listed in supplementary Table 1.

### Overview of molecular approach

Specimens were sampled non-destructively, by removing one or two legs, to preserve voucher integrity. Two sequencing strategies were applied depending on specimen age and preservation state. A total of 89 relatively fresh specimens were amplified and sequenced using the traditional Sanger method, whereas 288 historical specimens were processed using Oxford Nanopore Technologies (ONT) sequencing, optimised for degraded DNA from museum collections (Table 1).

### Sanger sequencing

Sequence data were generated from 76 out of 89 *Cytaea* specimens. Total genomic DNA was extracted from a single leg of each specimen using the DNeasy Blood and Tissue Kit (Qiagen, Hilden, Germany), following the manufacturer’s protocol, with the exception of the lysis step, during which samples were incubated for approximately 72 hours to improve yield from lightly degraded material. Amplification was carried out using the primers bcdF01 and bcdR04 (Dabert *et al*., 2010). PCR reactions were performed in 10 µL volumes containing 5 µL of Type-it Microsatellite PCR Master Mix (Qiagen), each primer at a final concentration of 0.5 µM, and 4 µL of DNA template. Thermocycling conditions included an initial denaturation at 95°C for 5 minutes, followed by 35 cycles of 30 seconds at 95°C, 60 seconds at 50°C, and 1 minute at 72°C, with a final extension step at 72°C for 5 minutes. PCR products were visualised on a 1% agarose gel to verify amplification success. Samples yielding clear bands were enzymatically purified using Exonuclease I and FastAP (Thermo Fisher Scientific, USA), and bidirectionally sequenced using BigDye Terminator v3.1 kit on an ABI Prism 3130xl Genetic Analyzer (Applied Biosystems), according to the manufacturer’s instructions. Sequence chromatograms were visually inspected for accuracy using FinchTV v1.3.1 (Geospiza Inc.). Sequences were aligned using MUSCLE (Edgar, 2004) under default parameters as implemented in Geneious v10.2.3 (Biomatters Ltd., New Zealand), and manually refined where necessary. The final dataset additionally includes 14 *Cytaea* sequences previously published by Trębicki et al. (2021).

### Mini-barcode primer design

To optimise COI barcoding for degraded DNA from historical museum specimens of *Cytaea*, we designed a set of primers targeting six short, overlapping (∼130 bp each) COI fragments (Table 2). Primer design was based on an alignment of 89 COI sequences previously obtained from fresh *Cytaea* specimens using Sanger sequencing. The external primers bcdF01 and bcdR04 (Dabert et al., 2010) were retained, while internal primers were adapted from Prosser et al. (2016) and modified to improve their match to *Cytaea* sequences while minimising unnecessary degeneracy. Several primer combinations and fragment lengths were tested on fresh *Cytaea* specimens with known full-length COI barcodes. The resulting amplicons were Sanger-sequenced by Macrogen Europe (Amsterdam, The Netherlands) following Querner et al. (2022), and the sequences were compared with the original barcodes to confirm their identity.

**Table 2.**
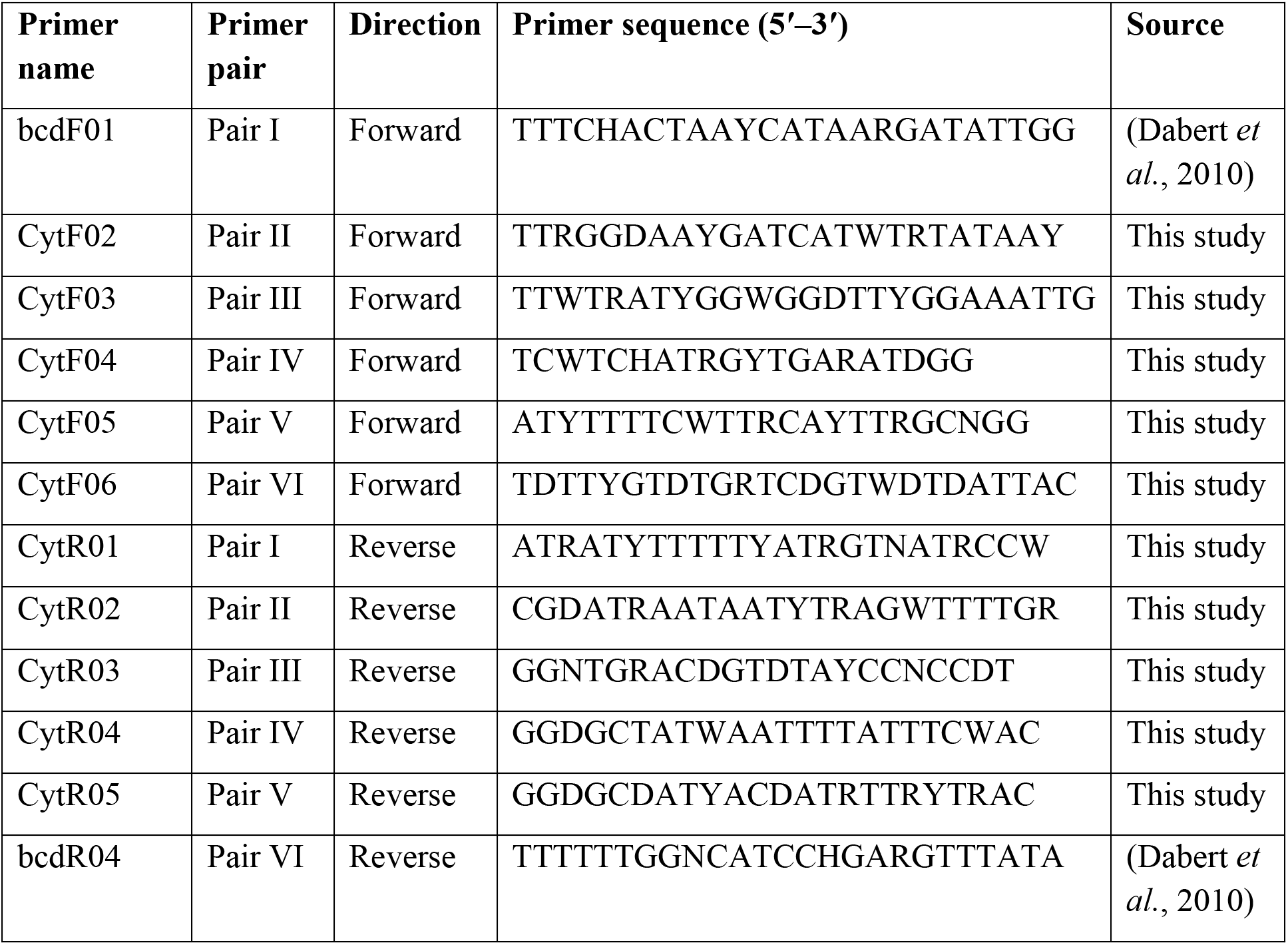
Primers used to amplify six overlapping COI mini-barcode fragments in *Cytaea*. Primer pairs are numbered I–VI. All primer sequences are presented in the 5′–3′ direction.

### DNA Extraction from Historical Specimens

Genomic DNA was extracted using the MagMAX CORE Nucleic Acid Purification Kit on a KingFisher Apex System (Thermo Fisher Scientific) to standardize processing and minimize manual handling. Lysis was performed overnight at 56°C, followed by approximately 30 minutes of automated magnetic bead purification. A total of 288 specimens were processed in a 96-well format across three independent extraction runs.

### Multiplex PCR of COI mini-barcodes

Each specimen underwent two multiplex PCR reactions to amplify six overlapping COI mini-barcodes (∼130 bp each), tailored for degraded DNA: Mix I: primers bcdF01–CytR01; CytF03–CytR03; CytF05–CytR05. Mix II: primers CytF02–CytR02; CytF04–CytR04; CytF06–bcdR04. Each PCR was performed in a total volume of 15 µL, consisting of: 7.5 µL of 2x QIAGEN Multiplex PCR Master Mix; 1.5 µL of primer mix (5 µM of each primer); 2 µL of DNA template; 4 µL of nuclease-free water. Thermal cycling conditions: initial denaturation at 95°C for 5 min; 40 cycles of: 95 °C for 30 s, 50 °C for 90 s, 72 °C for 1 min; Final extension at 68 °C for 30 min. PCR success was verified via 1% agarose gel electrophoresis.

### Indexing and library preparation

Amplicons from Mix I and Mix II were pooled separately for each specimen in a clean 96-well plate. The pooled amplicons were subjected to DNA repair and end preparation using the NEBNext® Ultra™ II End Prep Kit (New England Biolabs), following the ONT protocol. Amplicons were transferred to another 96-well plate and indexed using the ONT Native Barcoding Kit 96 V14 (SQK-NBD114.96). Native-barcoded libraries were pooled into one vial; cleanup was performed using AMPure XP beads at a bead-to-sample ratio (1:1) for expected short fragments. The cleaned library was quantified on the Qubit 4 Fluorometer with the High Sensitivity dsDNA Assay Kit (Thermo Fisher Scientific). Final preparation included adapter ligation using (SQK-NBD114.96) and NEBNext Quick Ligation Module (New England Biolabs). Pooled barcoded sample with Native Adapter (NA), Quick Ligation Reaction Buffer (5x) and T4 Ligase (NEBNext) was mixed and purified using AMPure XP beads with Short Fragment Buffer (SFB). DNA concentration was measured using the Qubit HS dsDNA kit. Libraries were diluted to the appropriate loading concentration for sequencing.

### Sequencing and bioinformatic processing

Sequencing was performed on a MinION device using Flongle flow cells (FLO-FLG001). Raw reads were basecalled and demultiplexed using Dorado versions 0.5.0–0.7.0 with the models dna_r10.4.1_e8.2_400bps_sup@v4.3.0 or dna_r10.4.1_e8.2_400bps_sup@v5.0.0. Cutadapt v4.9 was used to trim primers and assign reads to their corresponding amplicons. Initial consensus sequences were generated with abPOA from 100 randomly selected reads per amplicon per specimen. Consensus sequences were subsequently refined through iterative variant calling using FreeBayes, WhatsHap, BCFtools, and SAMtools, followed by polishing with Medaka, following an approach similar to that described by Prous et al. (2025). The processing script, including the software commands used, is provided as supplementary material 2. For future applications, the Minovar pipeline offers a better alternative for generating high-quality amplicon consensus sequences from complex mixtures (Tedersoo *et al*., 2026), although it was not used in this study.

### Mini-barcode validation and assembly

Consensus sequences for each of the six mini-barcodes were aligned to *Cytaea* COI reference sequences previously obtained via Sanger sequencing, checked using BLASTn against the NCBI nt database, and manually inspected to identify potential chimeras and non-target fragments. Chimeric, misidentified, and contaminant sequences were removed. Validated mini-barcodes from each specimen were assembled and aligned to the full-length COI barcode of a closely related *Cytaea* species obtained via Sanger sequencing. Missing mini-barcode regions were represented by Ns to preserve alignment structure. The sequences were aligned in Geneious 10.2.6 (Biomatters Ltd.) (Kearse *et al*., 2012), using MUSCLE (Edgar, 2004) with default parameters. The resulting alignment was edited and trimmed of primers, and checked for the absence of stop codons, double peaks (for sequences generated in Macrogen) and frameshifts using Geneious 10.2.6 (Biomatters Ltd.).

### Data availability

All newly generated sequences have been deposited in BOLD Systems. Associated specimen metadata (including collection data and voucher information) and comparative images of ethanol-preserved specimens are available within the corresponding BOLD records in a public dataset of DS-MUSEONT (https://doi.org/10.5883/DS-MUSEONT). Sequences of required quality were deposited in GenBank (to be provided) (https://www.ncbi.nlm.nih.gov/).

### Analysis of DNA degradation with specimen age

The relationship between specimen age and the number of successfully recovered nucleotides was analysed in Python 3.12 using the pandas, NumPy, Matplotlib, and SciPy libraries. To assess the strength and direction of the association between specimen age and nucleotide yield, a Spearman’s rank correlation test was applied using the spearmanr() function from scipy.stats. This non-parametric approach was chosen because the variables were not normally distributed and the relationship was expected to be monotonic rather than strictly linear. The Spearman’s ρ (rho) coefficient and corresponding p-value were calculated to determine the significance of the correlation. In addition, two regression models were fitted to explore the functional form of DNA degradation with age: (1) a linear regression model (y = a + b·x), representing a constant rate of nucleotide decline, and (2) a logarithmic regression model (y = a + b·log(x)), representing a rapid initial decrease that stabilizes over time. Both models were fitted using the linregress() function from the scipy.stats module. For each model, the regression equation and coefficient of determination (R²) were computed to assess goodness of fit.

### Geographic data and spatial analysis

Mapping and visualization of sampling localities were performed as follows. Coordinates were obtained from specimen labels or associated collection databases. Geographic visualization of sampling sites was performed in Python 3.12 using the libraries pandas, GeoPandas and Matplotlib. Sampling coordinates were converted into a GeoDataFrame using GeoPandas, and separate subsets were created for ONT-derived and Sanger-derived sequences. A basemap of country boundaries was obtained from the Natural Earth v5.0 dataset (1:110 million scale) and accessed programmatically via HTTPS. All spatial data were projected and visualized using standard GeoPandas workflows. Final map was generated for comparative visualization of geographic coverage between sequencing approaches.

### Diversity analysis

Sequences were clustered in BOLD Systems into molecular operational taxonomic units (OTUs) using the RESL algorithm (Ratnasingham and Hebert, 2013). Sequences of sufficient quality and length (>500 bp) were assigned to Barcode Index Numbers (BINs). Pairwise genetic distances among BINs were calculated using the Kimura 2-parameter (K2P) model (Kimura, 1980). Pairwise distance matrices and K2P heatmap visualizations were processed in Python 3.12 using NumPy, pandas, and Matplotlib.

### Phylogenetic backbone reconstruction

A reference phylogeny was reconstructed from 38 full-length COI sequences representing BIN-level diversity within *Cytaea*. Sequences were aligned using MAFFT with the E-INS-i algorithm and default parameters (Katoh and Standley, 2013), producing an alignment of 644 nucleotide positions with no missing data. Maximum likelihood (ML) inference was performed in RAxML-NG v1.2.2 (Kozlov *et al*., 2019) under the GTR+F+Γ4 substitution model, using 20 starting trees (10 random and 10 parsimony-based). The tree with the highest log-likelihood was retained, and node support was assessed using 300 nonparametric bootstrap replicates (Felsenstein, 1985). The resulting ML tree served as a fixed reference backbone for subsequent phylogenetic placement analyses.

### Phylogenetic placement and reliability assessment

A total of 185 ONT-derived COI mini-barcode sequences from historical specimens were aligned to the reference alignment using a profile-based approach and placed onto the ML backbone representing 38 BINs using EPA-ng v0.3.8 (Barbera *et al*., 2019). Placement used the GTR+F+Γ4 model and model parameters estimated for the backbone phylogeny. Results were exported in .jplace format and processed using GAPPA to extract placement statistics (Czech, Barbera and Stamatakis, 2020). Placement support was assessed using likelihood weight ratios (LWR) (Berger, Krompass and Stamatakis, 2011). The highest value (LWR_top1) quantified support for the best placement, while the difference between the two highest values (LWR_gap) measured separation from the next-best alternative. Based on LWR_gap, the two best placements were classified as well separated (LWR_gap ≥ 0.20), moderately separated (0.05 ≤ LWR_gap < 0.20), or poorly separated (LWR_gap < 0.05). Placements with LWR_top1 < 0.5 were additionally flagged as low-support. Support concentration and uncertainty were further assessed using the cumulative LWR of the three best placements (ΣLWR_top3) and Shannon entropy of the LWR distribution (Shannon, 1948).

## Results

### Reference dataset and full-length COI sequences

Sanger sequencing was attempted for 89 *Cytaea* specimens obtained from four natural history museum collections (Supplementary Table 1). The specimens were preserved in 75% ethanol and stored in museum collections for 2–10 years before sequencing. We obtained 71 complete COI sequences (>600 bp), 7 partial sequences (400–599 bp), and 11 failures (no COI data) (Table 3). The Sanger alignment comprised 644 nucleotide positions; translation of nucleotide sequences revealed no stop codons. Of these, 206 sites were variable, with a transition/transversion ratio R = 1.29.

**Table 3.** Results of Sanger-based COI barcoding of 89 *Cytaea* specimens from four natural history museum collections. The table presents specimen numbers, minimum and maximum time from collection to DNA extraction and sequencing (years), and sequencing outcomes classified as full-length COI barcodes (≥600 bp), partial barcodes (400–599 bp), or unsuccessful sequencing attempts.

| Museum | Specimens (n) | Age min | Age max | ≥600 bp | 400-599 bp | No COI sequence |
|---|---|---|---|---|---|---|
| QMB | 73 | 2 | 5 | 64 | 7 | 2 |
| USMB | 12 | 9 | 10 | 3 | 0 | 9 |
| MAGNT | 2 | 4 | 4 | 2 | 0 | 0 |
| WAMP | 2 | 2 | 7 | 2 | 0 | 0 |
| Total | 89 | 2 | 10 | 71 | 7 | 11 |

### Amplification success of COI mini-barcodes

We processed 288 *Cytaea* specimens across three 96-well plates, targeting six overlapping COI mini-barcode fragments per specimen. Across attempts (Seq I–III), overall success per mini-barcode (of 288 specimens) ranged from 31.60% (CytF05/CytR05) to 98.26% (CytF04/CytR04) (Table 4). Sequencing of longer fragments was limited: only 70 of 288 specimens (24.3%) yielded longer amplicons – approximately 300-500 bp – confirming the limited applicability of longer fragments in degraded museum DNA (Table 4).

**Table 4.**
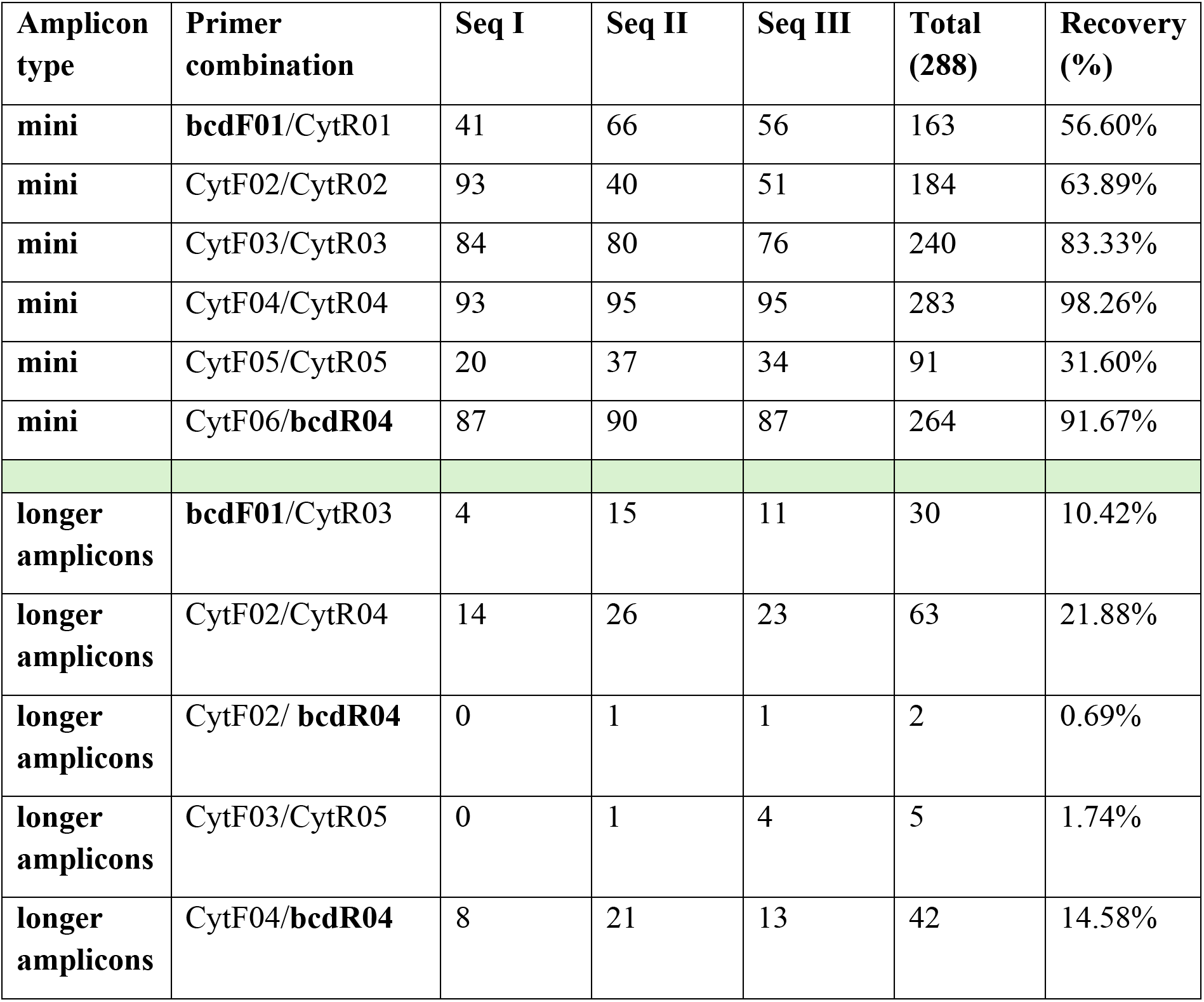
ONT-based barcoding results. Recovery of COI amplicons from 288 historical *Cytaea* specimens using ONT sequencing. Results are presented for three sequencing runs (Seq I–III), covering six overlapping mini-barcode fragments and five longer amplicons. Totals indicate the number of specimens yielding each amplicon, with percentages calculated relative to all 288 specimens.

### Recovery of COI sequences from museum specimens

The specimens represented a broad age range, from 13 to 141 years since collection (Table 1). COI sequences were successfully reconstructed for 284 of 288 specimens (98.6%), with lengths ranging from 115 to 663 bp (Figure 1). Among the 284 recovered sequences, 91 (32.0%) were 600–663 bp long, 64 (22.5%) were 500–599 bp, 105 (37.0%) were 300–499 bp, and 24 (8.5%) were 115–299 bp.

**Figure 1.**
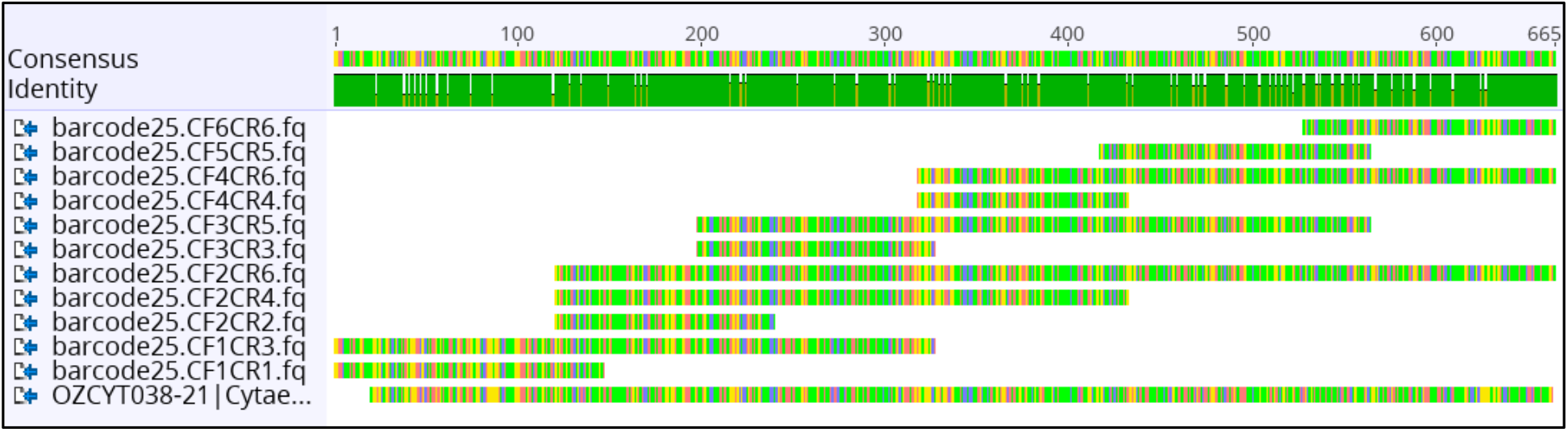
Example of COI barcode assembly from multiple overlapping mini-barcode fragments obtained from a single *Cytaea* specimen. A COI sequence from a closely related species was included as a reference to guide alignment and assembly.

The final alignment for species delimitation comprised 664 nucleotide positions for COI gene sequences representing 284 specimens. The nucleotide sequences were successfully translated into amino acid sequences without any stop codons. In the dataset, 258 nucleotide positions out of 664 were variable, and transition/transversion ratio (R) amounted to 1.25.

### Effect of specimen age on sequence recovery

Full-length COI barcodes were recovered from four specimens collected in 1949, 19 collected in 1971–1989, and 47 collected in 1992–2011. Specimen age was negatively correlated with the number of recovered nucleotides (Spearman’s ρ = −0.443, p = 2.79 × 10⁻¹⁵), with older specimens tending to yield shorter and less complete sequences. Under a linear approximation, recovered sequence length decreased by approximately 1.9 bp per year (slope = −1.93). The logarithmic model provided a better fit than the linear model (R² = 0.161 versus 0.114), suggesting a steeper decline in recovered sequence length among more recently collected specimens and a more gradual decline among older material (Figure 2).

**Figure 2.**
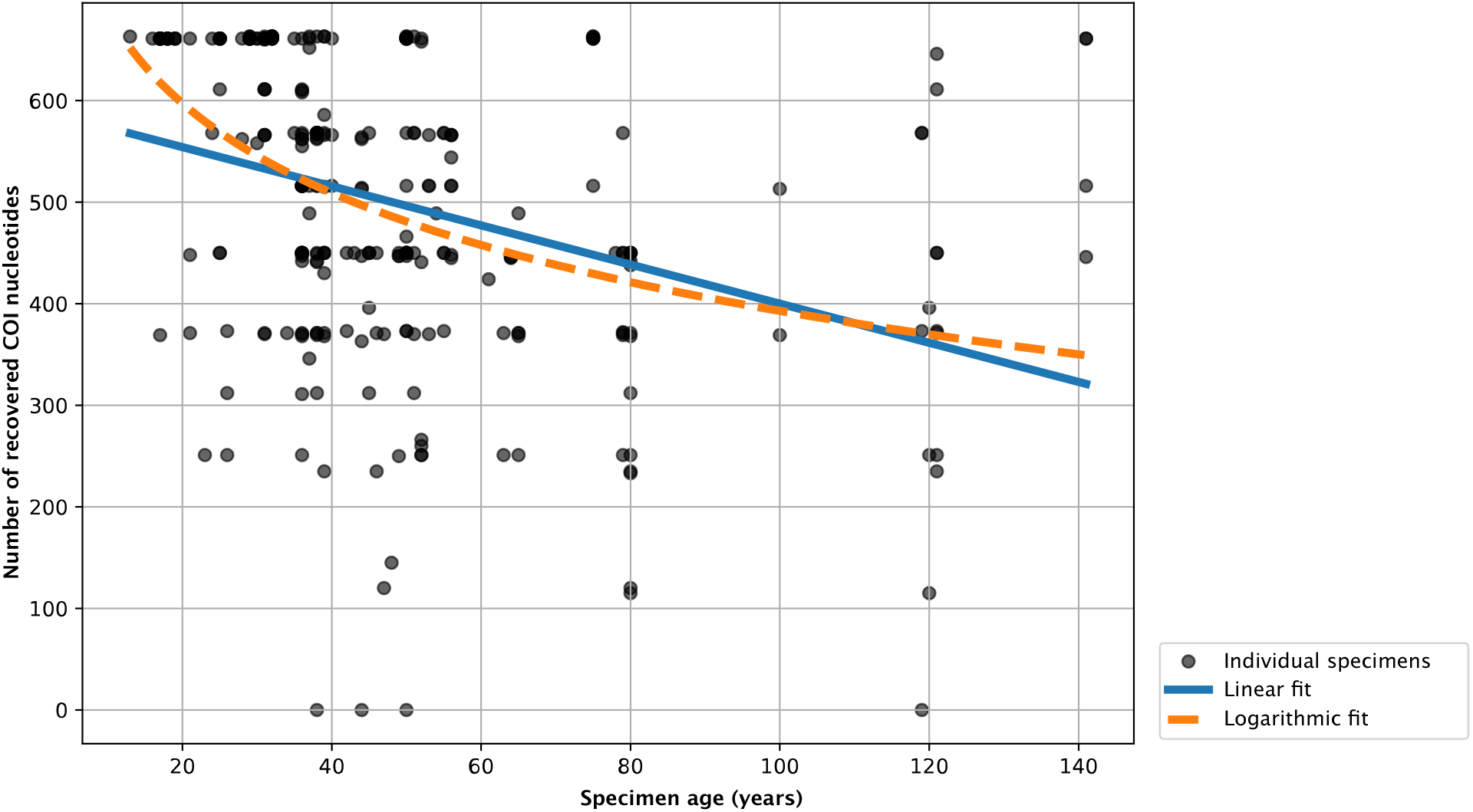
Relationship between specimen age and the number of recovered COI nucleotides. Each point represents one specimen. The four specimens from which no COI sequence was recovered are plotted at zero. The solid blue line shows the linear fit (y = 592.65 − 1.93 × age, R² = 0.114), and the dashed orange line shows the logarithmic fit (y = 977.71 − 126.99 × ln(age), R² = 0.161).

### Molecular diversity (OTUs/BINs)

Analysis of 78 Sanger-derived COI sequences using the RESL algorithm identified 16 OTUs, while 17 BINs were assigned in BOLD. The dataset of 284 ONT-derived COI sequences yielded 64 OTUs and 47 BINs. One large OTU contained 154 predominantly short sequences, including sequences assigned to multiple BINs, with a mean intra-OTU genetic distance of 6.95%. This indicates that the clustering approach merged distinct BIN-associated lineages within the fragmented dataset. The combined dataset of 362 COI sequences (78 Sanger-derived and 284 ONT-derived) yielded 65 OTUs and represented 54 BINs, of which 53 were newly contributed to BOLD. Pairwise K2P distances among BINs revealed generally strong genetic differentiation between most lineages, with the majority of comparisons exceeding commonly used species-level thresholds (Figure 3). However, several BIN pairs showed relatively low inter-lineage divergence, potentially representing either closely related species or intraspecific mitochondrial variation. Similarly, nearest-neighbour analyses demonstrated substantial variation in inter-BIN distances across the dataset (Figure 4). While many BINs were separated by >3% K2P divergence, several showed nearest-neighbour divergence close to or below 2%, indicating cases that may require additional taxonomic evaluation using integrative approaches.

**Figure 3.**
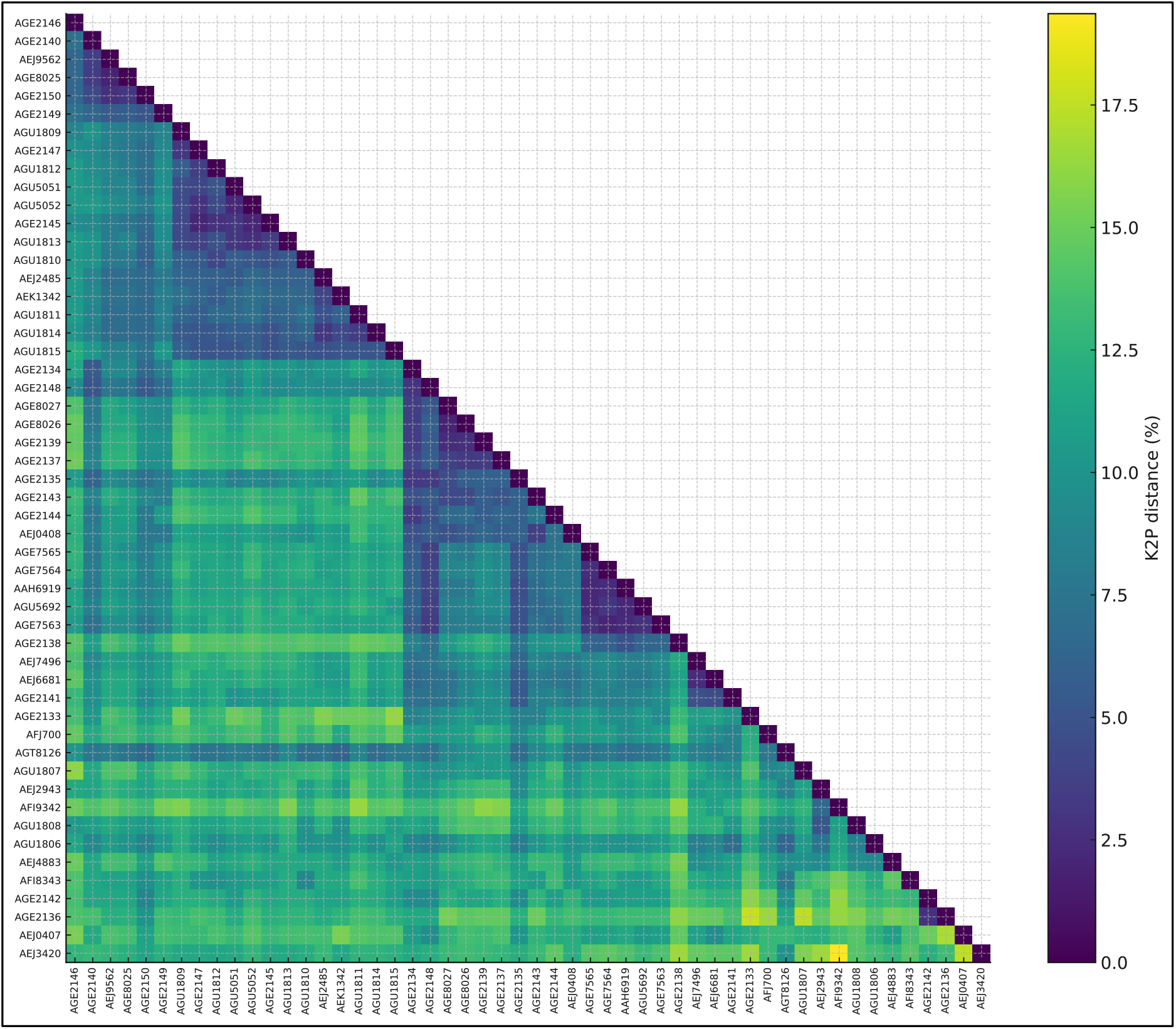
Pairwise K2P divergences among BINs (COI). Heatmap showing pairwise Kimura 2-parameter (K2P) sequence divergences between BIN-level groups. Row/column labels correspond to BIN identifiers; the matrix is symmetric and diagonal values equal zero by definition. Colour intensity follows the scale bar (0–20%), with lighter cells indicating higher divergence and darker cells indicating lower divergence. Values were calculated from COI sequences and are displayed in the input order without hierarchical reordering.

**Figure 4.**
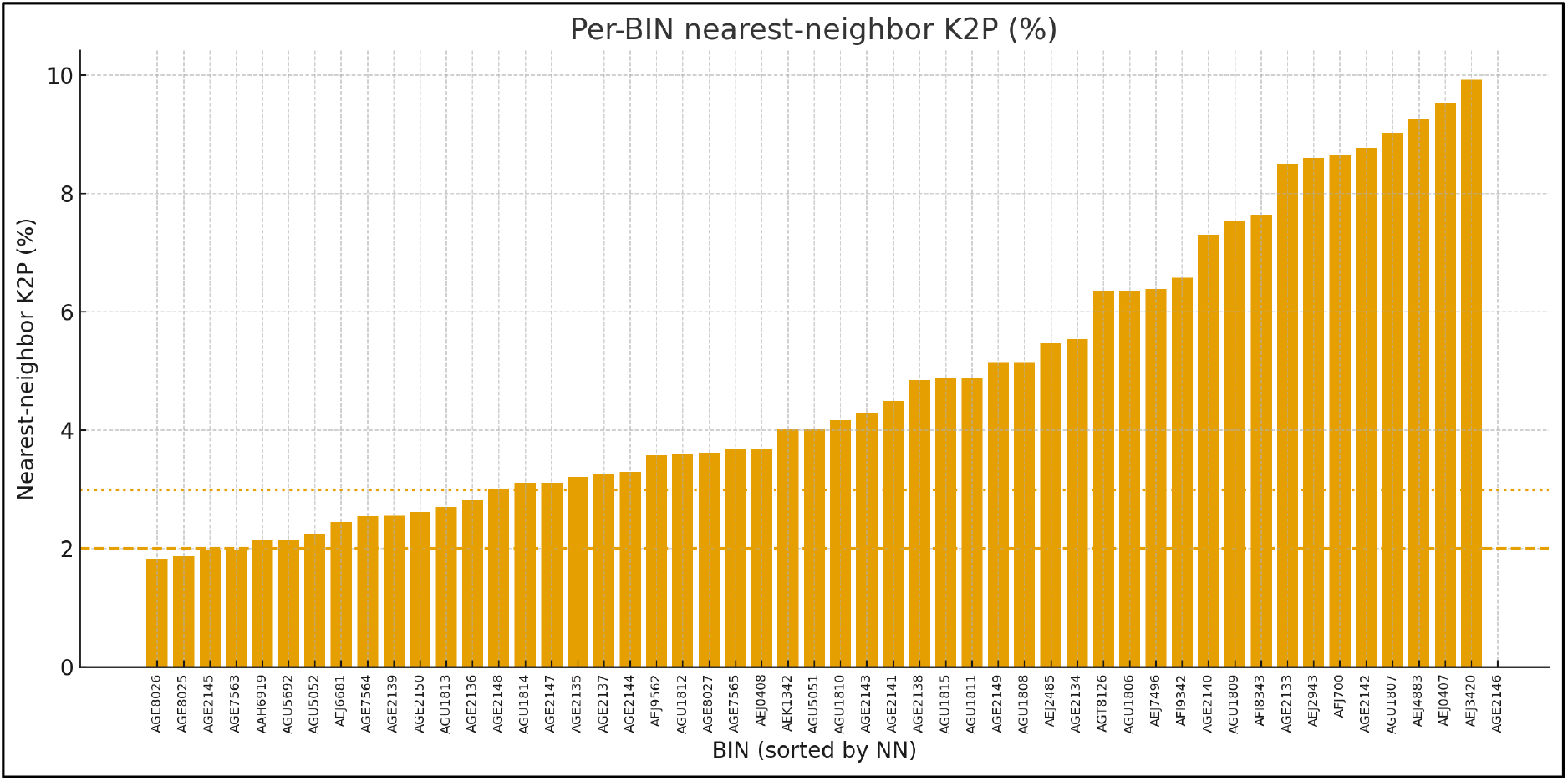
Nearest-neighbour K2P divergence per BIN (COI). Bar chart showing, for each BIN (x-axis; sorted ascending), the smallest pairwise K2P divergence to its nearest neighbouring BIN (y-axis). Dashed horizontal lines indicate reference thresholds at 2% and 3% sequence divergence. BINs above ∼3% show strong genetic differentiation, whereas BINs near or below ∼2% may represent closely related lineages requiring additional taxonomic assessment.

### Phylogenetic backbone reconstruction

The maximum likelihood phylogeny inferred from the COI alignment recovered several well-defined clades corresponding to the major species groups within *Cytaea* (Figure 5). The backbone dataset comprised 38 reference sequences representing BIN-level diversity across the genus. Several internal nodes received strong support (bootstrap ≥ 90), indicating stable clustering of BIN-level lineages despite limited sequence divergence. The topology revealed four main clades largely corresponding to the predefined species groups (*frontaligera*, *mitellata*, *alburna* and *sinuata*), although their monophyly received only moderate support at some deeper nodes. Within-clade relationships were generally better resolved, with several nodes receiving moderate to high bootstrap support. The *frontaligera* group formed a distinct lineage near the base of the tree and was represented by a small number of divergent sequences. The *sinuata* group constituted the largest and most structurally complex clade, containing multiple well-supported internal subclades. The *mitellata* group formed a relatively compact cluster with short internal branches, suggesting shallow divergence among its members. The *alburna* group formed a smaller but clearly delimited lineage within the tree. Several shallow nodes within the major clades exhibited moderate bootstrap support and short branch lengths. The alignment contained a high proportion of invariant sites (62.4%), which is typical for mitochondrial COI datasets and likely contributes to limited phylogenetic resolution among closely related BIN lineages. The resulting maximum likelihood topology was used as a fixed reference backbone for subsequent phylogenetic placement analyses of short COI mini-barcode sequences. Despite moderate support for deeper nodes, the backbone topology provided a stable framework for downstream placement analyses. Importantly, sequences generated using ONT consistently clustered together with their corresponding Sanger-derived reference lineages within the same BIN-level clades, without evident technology-specific segregation. This concordance between sequencing approaches supports the reliability of the museomics workflow and indicates that the ONT-derived consensus sequences retained sufficient phylogenetic signal for accurate placement within the *Cytaea* backbone topology.

**Figure 5.**
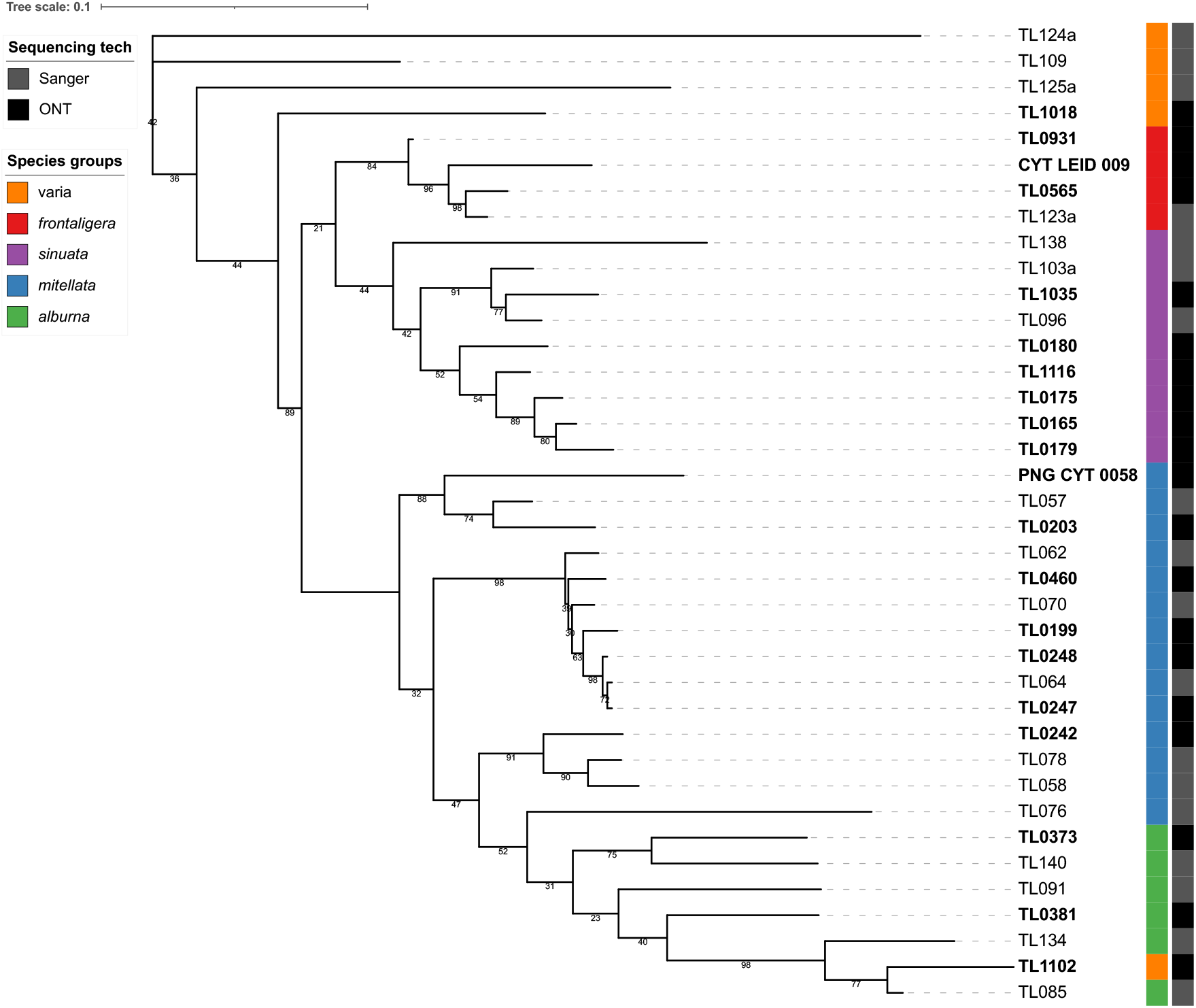
Maximum likelihood phylogeny of *Cytaea* based on full-length COI sequences. The tree was inferred using RAxML-NG from an alignment of 38 BIN-level reference sequences (644 bp). Numbers at nodes indicate bootstrap support values. Coloured bars indicate predefined species groups, while the adjacent column indicates sequencing technology used to obtain the sequence (Sanger or ONT). The tree served as the reference backbone for phylogenetic placement of short COI mini-barcode sequences obtained from historical specimens.

### Phylogenetic placement confidence

A total of 185 ONT-derived COI mini-barcode sequences were phylogenetically placed onto the reference backbone tree using EPA-ng. The distribution of maximum likelihood weight ratios (LWR) indicated generally high placement confidence across the dataset (Figure 6). Across all query sequences, the median maximum likelihood weight ratio (LWR_top1) was 0.86, with a mean of 0.76 and the upper quartile reaching 0.99. Overall, 87 sequences (47.0%) showed very high placement confidence (LWR ≥ 0.9), whereas 114 sequences (61.6%) had LWR values ≥ 0.7. In contrast, 42 sequences (22.7%) showed weak placement support (LWR < 0.5). The cumulative support of the three best placement positions (ΣLWR_top3) averaged 0.95, indicating that placement probability mass was typically concentrated on a small number of closely related branches. The distribution of LWR values was strongly skewed toward high-confidence placements, suggesting that short ONT-derived COI fragments frequently retained sufficient phylogenetic signal for reliable placement onto the reference tree. Placement uncertainty varied substantially among query sequences (Figure 7). Entropy values ranged from near zero for highly unambiguous placements to higher values indicating diffuse placement probability across multiple branches. As expected, maximum LWR values showed a strong inverse relationship with entropy, confirming that high-confidence placements correspond to strongly concentrated probability distributions across candidate branches, whereas ambiguous placements were associated with more evenly distributed placement probabilities.

**Figure 6.**
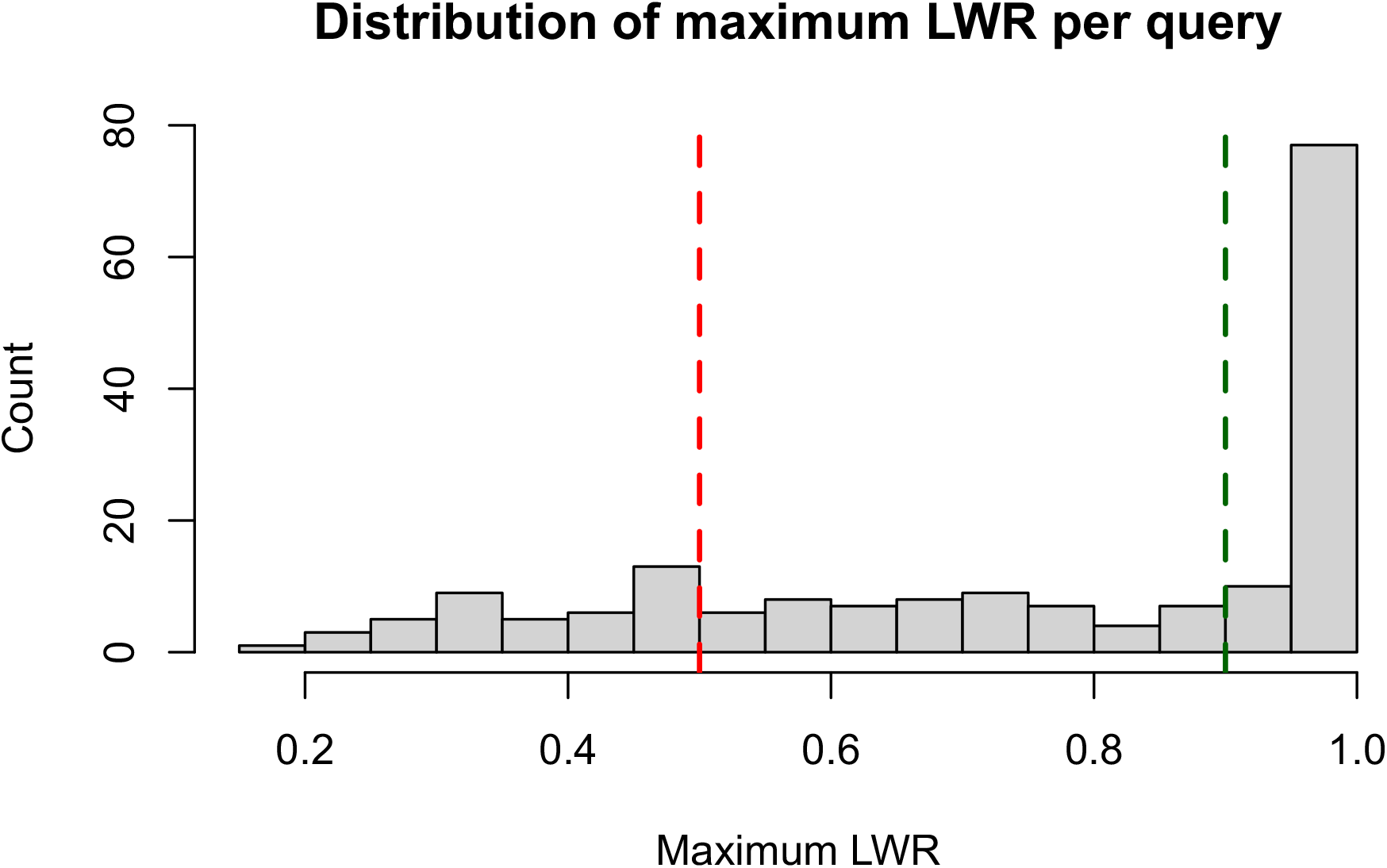
Distribution of maximum likelihood weight ratios (LWR_top1) for phylogenetic placements of ONT-derived COI mini-barcode sequences. The histogram shows the frequency distribution of the highest likelihood weight ratio obtained for each query sequence placed onto the reference backbone tree using EPA-ng. High LWR values indicate strong support for a single placement position, whereas lower values reflect increased ambiguity among alternative branches.

**Figure 7.**
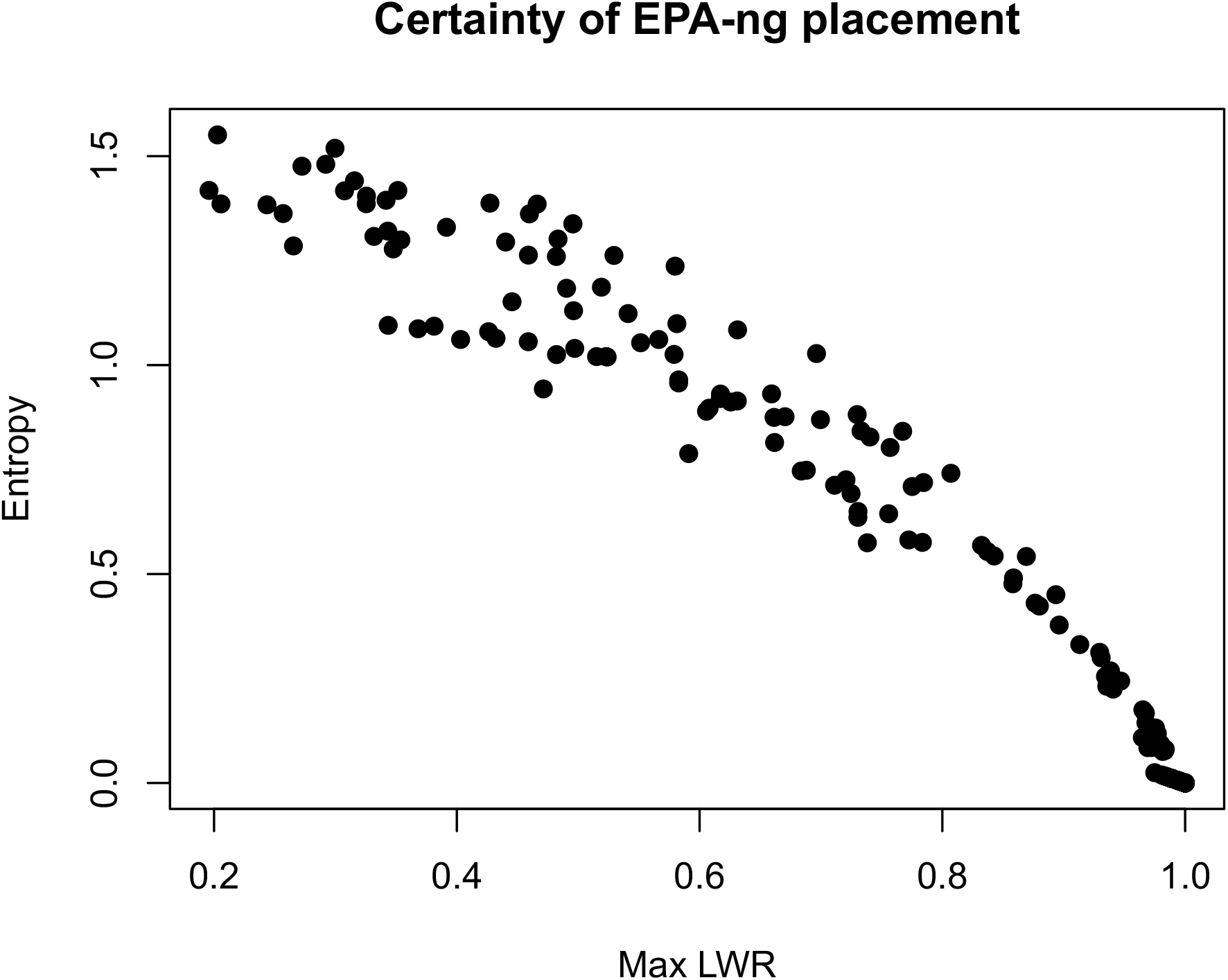
Relationship between maximum likelihood weight ratios (LWR_top1) and placement entropy for ONT-derived COI mini-barcode sequences. Each point represents a query sequence placed onto the reference backbone tree using EPA-ng. Lower entropy values indicate concentrated placement probability on a single branch, whereas higher entropy values reflect increased uncertainty among alternative placement positions.

### Placement patterns across the reference phylogeny

Phylogenetic placement of ONT-derived COI mini-barcode sequences onto the reference backbone tree revealed that most query sequences clustered within the major clades defined by the backbone phylogeny (Figure 8). Placements were broadly consistent with the species-group structure inferred from full-length COI sequences, indicating that short fragments retained sufficient phylogenetic signal for assignment to major lineages within the genus. The majority of query sequences were placed within the large clades of the sinuata and mitellata groups, which represent the most species-rich lineages in the dataset. Fewer placements occurred within the smaller alburna and frontaligera groups. In most cases, query sequences were placed close to their corresponding reference BIN representatives, forming local clusters within the backbone topology. The spatial distribution of placements across the tree suggests that the mini-barcode dataset captures a broad representation of the phylogenetic diversity present in the genus.

**Figure 8.**
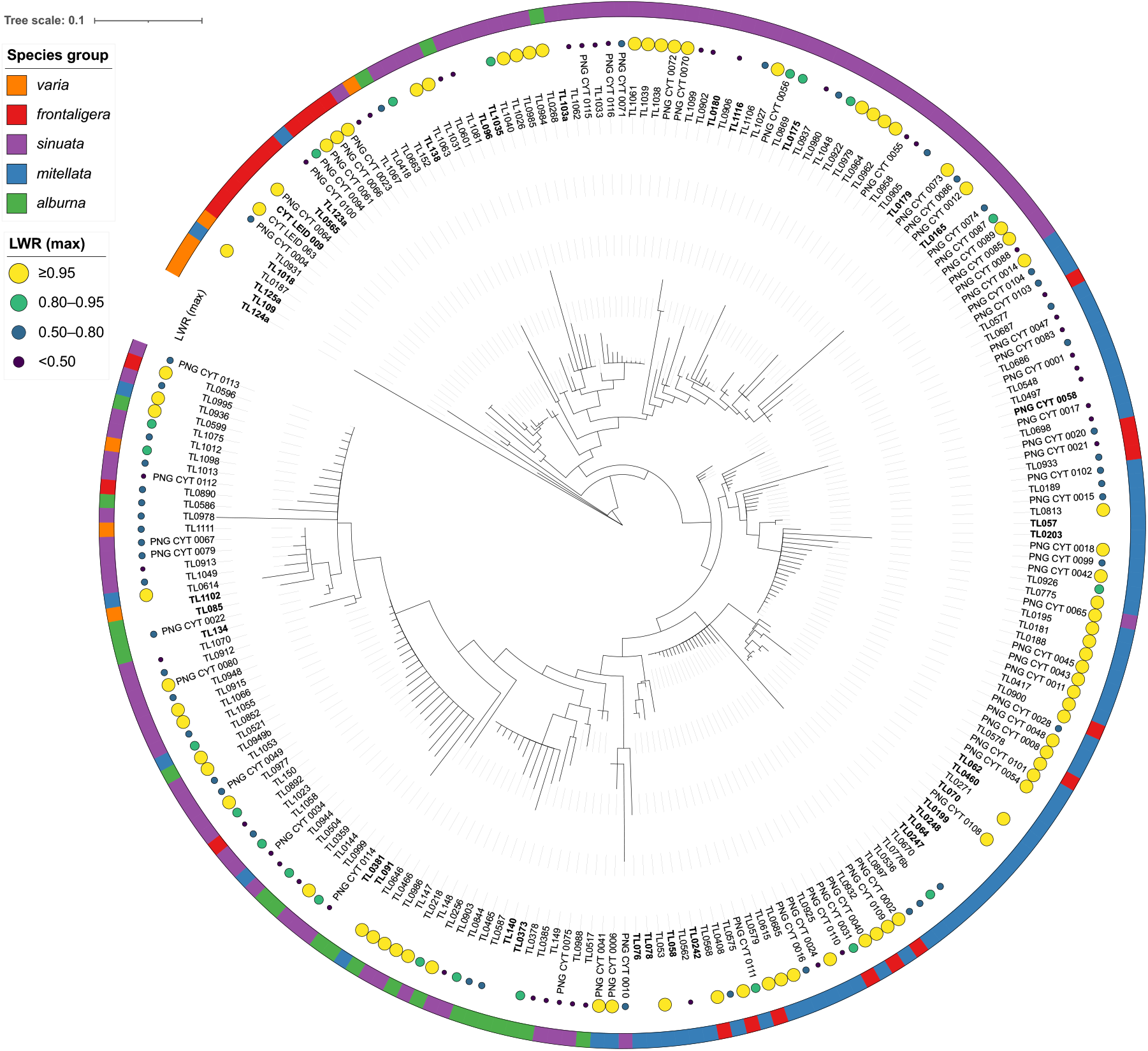
**Phylogenetic placement of ONT-derived COI mini-barcodes on the reference backbone tree of *Cytaea***. The backbone was inferred from 38 full-length COI reference sequences and held fixed during phylogenetic placement. The outer coloured ring indicates species-group assignments. Backbone reference labels are shown in bold. Circles mark only the query sequences placed using EPA-ng, and their colour and size indicate the highest likelihood weight ratio (LWR) among candidate placements for each query.

However, some placements were distributed across multiple neighbouring branches, indicating cases of ambiguous placement likely associated with limited phylogenetic signal in short COI fragments. Overall, the placement results demonstrate that short ONT-derived COI fragments can be reliably integrated into a reference phylogeny using evolutionary placement methods.

### Geographic coverage of sampling

Geographic distribution of sampled specimens revealed that the inclusion of ONT-derived sequences from historical museum material substantially expanded the spatial coverage of the dataset compared to Sanger-based sampling alone (Figure 9). While Sanger sequences were largely concentrated in more recently collected material, ONT sequencing enabled the incorporation of specimens from a broader range of localities, including regions and time periods otherwise underrepresented in the dataset. This resulted in a more comprehensive representation of the geographic distribution of *Cytaea* lineages.

**Figure 9.**
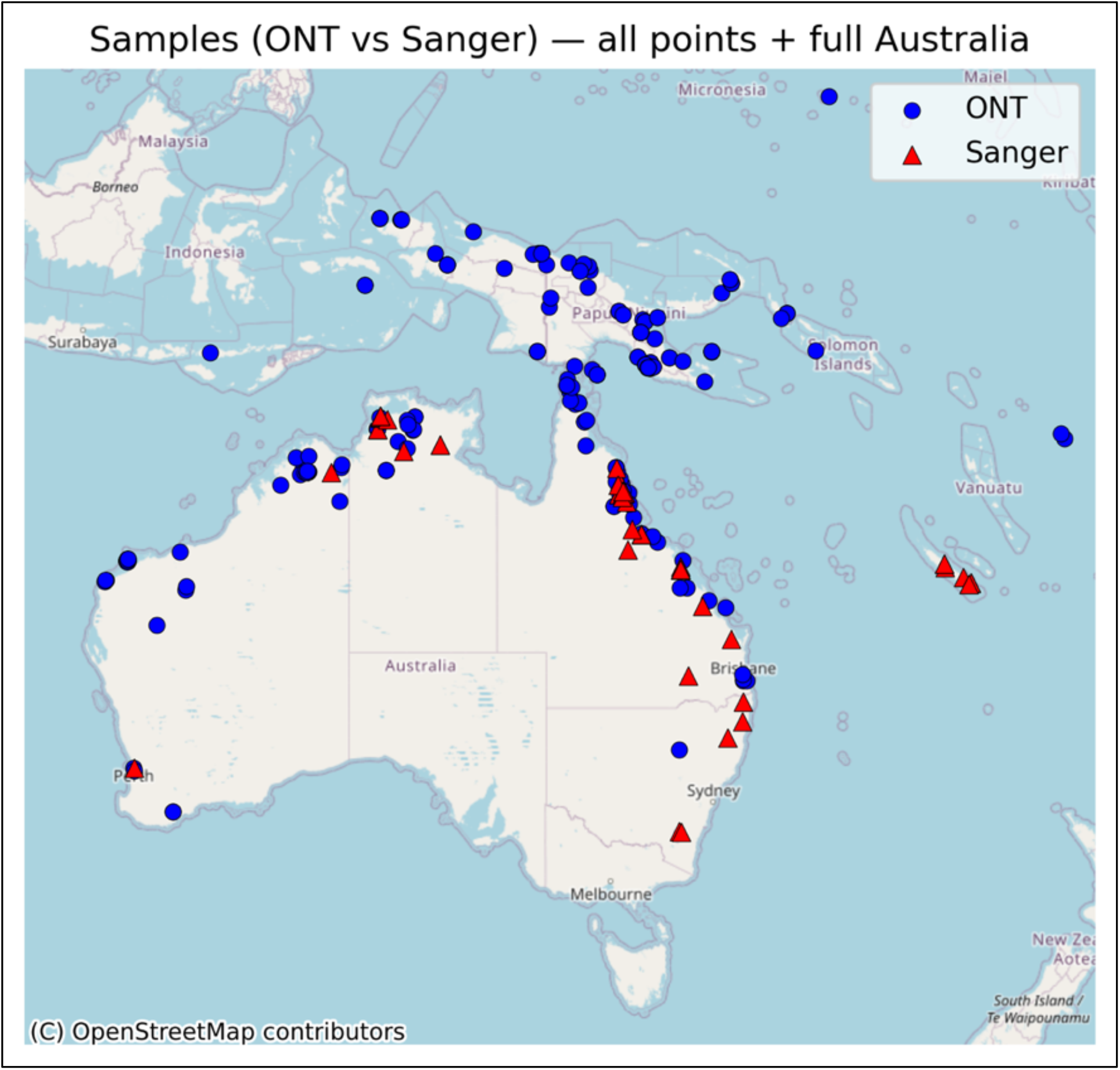
Sampling localities of fresh reference specimens sequenced using Sanger sequencing and historical museum specimens sequenced using ONT, illustrating the geographic coverage gained through inclusion of museum material.

## Discussion

We investigated DNA recovery from museum collections of the jumping spider genus *Cytaea*, for which fresh material was largely unavailable and historical specimens represented the primary source of genetic data. Standard DNA barcoding approaches proved largely ineffective, mainly due to repeated failures in amplifying COI from degraded DNA, necessitating the development of an alternative strategy tailored to highly fragmented templates. Consistent with previous studies, these challenges reflect the limitations of museum-derived DNA, including degradation, amplification bias, and variable sequencing success (Raxworthy and Smith, 2021; Ferrari *et al*., 2023; Fong *et al*., 2023). Despite these constraints, our ONT-based workflow recovered COI data from 284 of 288 specimens (98.6%), including specimens more than a century old, without repeated amplification or sequencing attempts within the final workflow. Of the 284 reconstructed sequences, 260 (91.5%) contained at least 300 bp of resolved sequence. These results demonstrate the potential for cost-efficient and scalable barcoding of historical collections, providing a practical framework for integrating museum specimens into modern molecular studies.

### Protocol development and practical advantages

Initial attempts to obtain COI barcodes from museum specimens often resulted in poor-quality sequences or amplification of non-target DNA, including bacterial or fungal sequences, even in relatively recently collected specimens (∼7 years old). This suggests that specimen age is not the sole factor determining barcode recovery success, consistent with previous reports of variable DNA preservation in ethanol-preserved spiders (Vink *et al*., 2005; Miller *et al*., 2013). The improved recovery achieved here reflects the combined use of short, overlapping amplicons, multiplex PCR, and ONT sequencing, following the principle of reconstructing COI barcodes from short fragments (Prosser *et al*., 2016). A practical advantage is that multiplexed amplicons can be sequenced together and target reads distinguished from non-target reads, whereas a Sanger-based approach would typically require separate sequencing reactions for individual fragments. This can reduce laboratory handling when processing multiple fragments per specimen. Previous studies have also demonstrated the scalability and cost-efficiency of ONT barcoding, although costs depend on sample number and workflow design (Srivathsan *et al*., 2019, 2021, 2024; Cuber *et al*., 2023; Hebert *et al*., 2025). ONT platforms offer flexibility for laboratories processing museum specimens locally, with lower-capacity Flongle flow cells suited to smaller batches and standard MinION flow cells accommodating larger batches. This is useful for taxonomic projects in which specimens become available gradually. External sequencing services may impose minimum batch sizes or additional charges for small submissions, while recovery from degraded material remains uncertain. Local processing gives researchers greater control over batch size, troubleshooting, and costs.

### DNA degradation and effects of specimen preservation

Museum spider specimens are preserved under a variety of conditions, and DNA preservation depends on multiple factors, including storage temperature, preservative type and concentration, and specimen history. High-concentration ethanol combined with low-temperature storage generally favours DNA preservation but may increase specimen brittleness, creating a trade-off between molecular preservation and morphological examination (Vink *et al*., 2005; Marquina *et al*., 2021). Desiccated specimens may exhibit tissue damage and highly degraded DNA, and rehydration does not necessarily restore their original morphological condition (Shikak, Anderegg and Cushing, 2025). Our *Cytaea* specimens originated from several museums and were primarily preserved in approximately 75% ethanol, with heterogeneous and incompletely documented storage histories. Some material was collected in tropical regions and may have been exposed to elevated temperatures before reaching the collections. Some older specimens yielded more complete sequences than younger material, suggesting that age alone did not determine recovery. This is consistent with findings that specimen age may affect DNA fragment length more strongly than total DNA yield or endogenous DNA content (Djernæs *et al*., 2026). Preservation history may therefore explain part of the variation observed here, although its individual effects could not be assessed. For newly collected spiders, a practical compromise is to preserve one or two detached legs separately in 96% ethanol at low temperature for molecular analyses, while retaining the remaining specimen in approximately 75% ethanol for morphological study. In our experience, short-term transfer from high- to lower-concentration ethanol does not necessarily restore appendage flexibility, and legs may detach during examination. Additional, potentially time-consuming softening procedures may therefore be required.

### DNA extraction strategies and lysis efficiency

DNA recovered from historical museum specimens is often highly degraded and fragmented, making tissue lysis and DNA extraction critical steps in molecular analyses. An effective extraction protocol should maximise DNA recovery while minimising further fragmentation and loss of genetic material. In our preliminary tests with historical *Cytaea* specimens, magnetic bead-based extraction yielded more usable DNA than the phenol–chloroform and silica column-based methods tested, although no systematic comparison was performed. Different outcomes reported for mammalian specimens emphasise that extraction performance depends on the material and preservation conditions rather than a universally superior method (Molbert *et al*., 2023). Our observations therefore support adapting established protocols to the specimens under study (Patzold, Zilli and Hundsdoerfer, 2020; Djernæs *et al*., 2026). The selected protocol achieved nearly complete soft-tissue digestion while preserving the external morphology of sampled legs, whereas other preliminary treatments frequently left residual tissue, potentially limiting DNA recovery. Our protocol used overnight incubation at 56°C, although alternative lysis conditions warrant further testing. Prolonged incubation at 42°C improved DNA recovery from historical insects, providing a potential direction for optimisation in spiders (Djernæs *et al*., 2026). Such optimization should consider DNA recovery, the risk of additional degradation, and preservation of diagnostic morphological structures (Tang *et al*., 2026).

### PCR bias and primer-specific performance

Amplification success varied markedly among primer pairs, from 31.6% for CytF05/CytR05 to 98.3% for CytF04/CytR04, indicating uneven recovery of COI regions. Sequence variability at internal primer-binding sites can complicate the design of multiple closely spaced primer pairs and limit their universality, even among closely related taxa (Prosser *et al*., 2016; Schäffer, Zachos and Koblmüller, 2017). Primer-template mismatches, sequence composition, and secondary structure may therefore contribute to differences in recovery, although their contributions were not tested here. Further optimisation could focus on poorly performing primer pairs and PCR conditions. Additives such as BSA or DMSO may also improve amplification under particular conditions (Farell and Alexandre, 2012), but their effects require validation in this multiplex system.

### Fragment length distribution and recovery patterns

Despite differences in amplification success among primer pairs, the use of multiple overlapping fragments enabled the reconstruction of relatively long COI barcodes from many historical specimens. Of the recovered sequences, 32.0% reached 600–663 bp, 59.5% were 300–599 bp long, and only 8.5% were shorter than 300 bp. By comparison, longer individual PCR amplicons (∼300–500 bp) were successfully recovered from only 70 of 288 specimens (24.3%), highlighting the value of targeting short, overlapping mini-barcodes for COI recovery from historical material. This approach has previously been shown to enable the recovery of substantial portions of COI when amplification of longer fragments is unsuccessful (Shokralla *et al*., 2011; Hebert *et al*., 2013; Prosser *et al*., 2016). Even shorter sequences may provide useful information for species identification, although their discriminatory power depends on the COI region and taxonomic group examined (Meusnier *et al*., 2008; Hernández-Triana *et al*., 2014; Yeo, Srivathsan and Meier, 2020).

### Effect of specimen age on sequence recovery

Although the assembly of overlapping mini-barcodes enabled substantial COI recovery from historical specimens, sequence completeness generally decreased with specimen age. Nevertheless, COI sequences were recovered from specimens up to 141 years old. The slightly better fit of the logarithmic model (R² = 0.161 versus 0.114 for the linear model) is consistent with a steeper decline in recovered barcode length among younger specimens, followed by a more gradual decrease among older specimens. The age-related decline in recovered COI barcode length is consistent with the trend towards shorter DNA fragments in older herbarium specimens (Porrelli *et al*., 2026). However, both models explained only a limited proportion of the observed variation, and the estimated linear decline of approximately 2 bp per year describes recovered barcode length rather than a directly measured rate of DNA degradation. These findings highlight that specimen age is only one of several factors influencing barcode recovery from historical material (Hebert *et al*., 2013; Djernæs *et al*., 2026).

### Incomplete barcode data and analytical challenges

The recovery of incomplete COI barcodes also presents challenges for downstream analyses. Mini-barcode datasets require careful validation of individual fragments and assembled sequences to identify non-target amplification and potential chimeras. Shorter, reliable sequences are preferable to longer assemblies containing questionable regions. Partial barcodes may also lack sufficient overlap for informative distance comparisons. In addition, sequences shorter than 500 bp do not meet the minimum length requirement for generating new BINs in the BOLD system (Ratnasingham and Hebert, 2013). These limitations were reflected in our ONT dataset, which yielded 64 OTUs but only 47 BINs. Notably, one OTU contained 154 sequences, including representatives assigned to different BINs, with a mean intra-OTU genetic divergence of 6.95%. This suggests that clustering of short and incomplete COI sequences may fail to distinguish genetically divergent lineages, highlighting the importance of complementary approaches such as phylogenetic placement and morphological examination.

### Backbone-based phylogenetic placement

Given the limitations of clustering incomplete COI sequences, phylogenetic placement provides a complementary approach for interpreting museum-derived mini-barcodes. We used EPA-ng to place short COI sequences onto a reference ML tree representing 38 BINs (Berger, Krompass and Stamatakis, 2011; Barbera *et al*., 2019). This approach allowed incomplete sequences to be analysed within a common phylogenetic framework without reconstructing relationships from highly incomplete alignments. The median likelihood weight ratio of the best placement (LWR_top1) was 0.86, with 47% of query sequences reaching values ≥ 0.9, indicating strong relative support for many placements. However, placement reliability depends on the phylogenetic information retained in short sequences and the quality and taxonomic coverage of the reference tree. Importantly, high LWR values do not necessarily indicate correct species identification, as placements remain conditional on the available reference sequences. Further analyses comparing individual COI mini-barcode regions could help determine which fragments provide the most reliable phylogenetic placements.

### Comparison with alternative genomic approaches

Although phylogenetic placement helps interpret incomplete COI sequences, our workflow recovers only a single mitochondrial locus, limiting its capacity to resolve deeper relationships and distinguish species histories from mitochondrial gene histories. Genome skimming, target capture, RADseq, and whole-genome sequencing provide broader genomic data and greater potential for resolving phylogenetic and evolutionary relationships, but generally involve higher costs and more complex laboratory and analytical workflows (Derkarabetian, Benavides and Giribet, 2019; Wahlberg, 2025; Lumen *et al*., 2026). For museum genome skims, workflows are available for batch assembly and annotation of mitochondrial and nuclear ribosomal sequences (White *et al*., 2025) and for recovery and validation of protein-coding DNA barcodes (Parsons, Vos and Price, 2026). Targeted COI mini-barcoding offers a simpler alternative where barcode recovery and taxonomic screening of historical material are the primary objectives (Folly *et al*., 2025). Such screening can identify genetically distinct lineages and guide representative sampling for subsequent genomic analyses, including UCE capture, making these approaches complementary rather than interchangeable.

### Taxonomic implications and biological insights

Museum-derived COI sequences expanded the geographic coverage and barcode diversity represented in our *Cytaea* dataset, with 37 of the 54 BINs represented exclusively by historical specimens sequenced using ONT. Museum collections provided access to geographically dispersed material accumulated over decades, including localities poorly represented among the available fresh specimens. Their inclusion therefore provides a broader basis for investigating lineage diversity and distributions within the genus. Where barcode libraries lack links to reliably identified historical specimens and name-bearing types, molecular clusters may remain disconnected from formally described species. Museum mini-barcoding can help establish these links and improve the taxonomic interpretation of barcode diversity (Letsch *et al*., 2025). This is particularly valuable in *Cytaea*, where many nominal species remain poorly documented and diagnostic morphological characters can be subtle, quantitative, and affected by intraspecific variation, sexual dimorphism, or preservation state. Recovering COI from historical reference material can therefore support both the interpretation of molecular clusters and the reassessment of earlier identifications. The additional BINs identify lineages requiring further taxonomic investigation and may include overlooked species. However, BINs should not be equated directly with species boundaries without supporting evidence. Museum mini-barcoding thus provides a practical foundation for an integrative revision of *Cytaea*, allowing candidate lineages to be linked to vouchers and prioritised for detailed morphological and molecular examination.

### Future directions and methodological perspectives

Further optimisation of museum mini-barcoding workflows could reduce costs, simplify laboratory procedures, and expand their applicability to other taxonomic groups. Such improvements should address the entire workflow, from DNA extraction and amplification to sample indexing and sequence validation, rather than sequencing technology alone (Ferrari *et al*., 2023). DNA extraction was automated in this study primarily for processing efficiency and laboratory logistics. Adapting magnetic bead purification to a manual format could reduce equipment requirements and facilitate implementation in laboratories equipped for routine molecular work. Low-cost magnetic stands, including 3D-printed holders fitted with neodymium magnets, could support manual purification without an automated platform. In previous work on Madagascan huntsman spiders (Sparassidae), we adapted the magnetic bead-based DNA extraction protocol of Tin et al. (2014) for manual use with 3D-printed magnetic racks, avoiding the need for extraction kits and plasticware specific to automated platforms (Uharov *et al*., 2026). Published low-cost museum DNA extraction protocols provide further options for adaptation and validation (Hall *et al*., 2023; Holmquist *et al*., 2025). Further savings could be achieved through sample-specific primer tags as an alternative to the ONT Native Barcodes used here (Srivathsan *et al*., 2019, 2021, 2024; Hebert *et al*., 2025). Their implementation in multiplex mini-barcoding would require validation of sample demultiplexing, amplification balance, and sequence recovery. The approach could also be adapted to group-specific primers and other markers, including mitochondrial 16S and nuclear 18S and 28S rRNA genes. Marker selection should depend on the taxonomic group, research question, and phylogenetic scale of interest.

## Conclusions

Our study demonstrates that ONT-based mini-barcoding enables efficient recovery of COI sequences from historical museum specimens, including material more than a century old. By combining magnetic bead-based DNA extraction, multiplex amplification of short overlapping COI fragments, and ONT sequencing, we recovered COI data from 284 of 288 *Cytaea* specimens (98.6%), substantially expanding the barcode diversity represented in the genus. Fragmented barcodes require rigorous validation and may complicate clustering-based analyses, while backbone-based phylogenetic placement provides a complementary framework for their interpretation. Overall, this workflow offers a cost-effective and scalable approach to integrating historical specimens into barcode reference libraries and helping to link molecular lineages, museum vouchers, and, where available, name-bearing types. The use of a compact sequencer, together with the potential to adapt DNA extraction for manual processing, could make the workflow accessible to smaller laboratories with limited equipment budgets.

## Supporting information

Supplementary Table 1. Specimen metadata and BOLD sequence records.

